# Intra- and interspecific variations in genome sizes of *Agaricia* corals from Curaçao

**DOI:** 10.1101/2023.08.23.554453

**Authors:** Dina Mae L. Rañises, Maria Juliana Vanegas Gonzalez, Mohammed M. Tawfeeq, Florence Rodriguez Gaudray, Maria Celia (Machel) D. Malay, Mark Vermeij, Jean-François Flot

## Abstract

Genome size is a fundamental biological trait that is known to exhibit high diversity among eukaryotic species, but its intraspecific diversity has only scarcely been studied to date. In scleractinian corals, genome size data are only available for a few species. In this study, intra- and interspecific variations in genome size of the coral genus *Agaricia* collected from Curaçao were investigated. Morphology was congruent with genetic analyses of the nuclear markers internal transcribed spacer 2 (ITS2) and L-threonine 3-dehydrogenase (TDH) in delimiting three *Agaricia* species among our samples. A refined Feulgen Image Analysis Densitometry (FIAD) protocol yielded genome sizes that ranged from 0.359 pg to 0.593 pg within this genus (a 1.7-fold range). The highest intraspecific variation in genome size was recorded in the depth-generalist *A. lamarcki* (1.5-fold range), followed by the depth specialist *A. humilis* (1.4-fold range) and *A. agaricites* (1.3-fold range), the species with an intermediate depth distribution. The mean genome size of *A. agaricites* (0.495 pg) was significantly larger than that of *A. lamarcki* (0.448 pg) and *A. humilis* (0.434 pg). No correlation between average genome size and nucleotide polymorphism *π* was detected, but we found an almost linear correlation between intraspecific variance of genome size and *π* of ITS2 (Pearson’s r = 0.984, p = 0.113). Genome size and collection depths of both *A. lamarcki* (Pearson’s r = 0.328, p = 0.058) and *A. agaricites* (Pearson’s r = -0.270, p = 0.221) were also not significantly associated. To our knowledge, this study provides the first account of intraspecific variation in corals; the apparent correlation detected between the nucleotide polymorphism of a species and the variance of its genome size will have to be tested using a larger taxonomic spectrum of scleractinian corals as well as in other groups of animals.

## Introduction

The genome size of an organism, commonly referred to as the “C-value”, is broadly defined as half of the amount of DNA in a somatic nucleus (1). It is highly variable among organisms (2), spanning a more than 200,000-fold variation among eukaryotes (3) and a 7,000-fold variation among animals (4). Such variation has been suggested to result from polyploidization events, recombination events, differential gain or loss of genes, and variable amounts of non-coding DNA (e.g transposable elements) in the genome (5–9).

The hypotheses underlying the variation in noncoding DNA among genomes could be summarized into two main categories: adaptive and non-adaptive theories (10). Adaptive theories suggest that variation in noncoding DNA has significant phenotypic effects linked to the fitness of an organism, and is thus under the control of natural selection. However, there has been no single pattern observed in previous studies that have investigated the relationship between genome size and adaptability or ecological tolerances of species (11, 12). In rayfinned fishes, Smith & Gregory (2009) (6) did not observe any relationship between genome size and their Red List status. Smaller genomes in plants are associated with higher adaptability and invasiveness (13), while larger genomes in salamanders were reported to possibly constrain species from occupying different larval habitats (14). A contrasting pattern was reported by dos Santos et al. (15) who found a significant positive association between genome size and ranges of temperature and salinity (considered as proxies of niche breadth) in reef fishes, suggesting that species with larger genomes tend to tolerate a wider range of environmental conditions. Non-adaptive theories postulate that mutation and genetic drift are the main forces underlying genome size variation (16). The passive accumulation of slightly deleterious mutations that would otherwise be removed by efficient selection is associated with a low effective population size (Ne), which results in stronger genetic drift. Lefébure et al. (2017) (10) have reported a negative correlation between genome size and effective population size, where populations with low Ne are expected to have larger genome sizes because slightly deleterious increases in genome sizes are suggested to be less efficiently removed by purifying selection (10, 16). The relative contributions of these evolutionary processes to genome size variation still remain to be explored. Nevertheless, the genomic sequences linked to this variation are the targets of different evolutionary pressures that influence the potential evolutionary fates of species (17). Despite this, genome size is recorded in only 6,222 animals in the Animal Genome Size Database, among which 3,793 vertebrates (i.e., 61%) (18). The extent to which genome size varies within species is also not well-documented and may be more common than what is currently reported in a few studies (19–22).

Scleractinian corals, the major group of reef-building organisms, consists of around 1, 676 accepted species (23) that are increasingly threatened by anthropogenic pressures (e.g increased sedimentation, pollution, overexploitation) (24) and global factors associated with climate change (e.g mass coral bleaching events and reduced calcification rates due to ocean acidification) (25). These continuing threats to coral reefs demand a deeper understanding of evolutionary traits linked to their adaptive potential. Certain evolutionary traits have already been suggested to influence the extinction risk of corals, including the presence of symbionts, resistance to bleaching, small or solitary colonies, and vertical and global distribution (26). Other traits, such as genome size, have almost not been studied. Only the genome sizes of 6 scleractinian coral species have been measured directly, whereas genome size estimates from sequence data are available for 23 additional species (see Supplementary Taable 1).

Coral species that are found across both the shallow (<30 m) to mesophotic (30-150 m) depths (‘depth-generalists’) are proposed to be likely candidates for the deep-reef refuge hypothesis (27), which postulates that the environmentally more stable mesophotic zone could potentially serve as a local source of propagules to replenish the shallow water population after extreme catastrophic events. Deep-water environments are assumed to be more stable in terms of environmental and climatological stressors compared to shallow reef communities (28, 29). If mesophotic coral populations are less subjected to regular stressors than their shallow-water counterparts, then the relaxation of selection on them could lead to the accumulation of transposable elements and thus to larger genome sizes (9). This hypothesis, however, remains to be tested in corals. Investigating genome size variation between depth-generalists and depth-specialists could also potentially answer the question of whether variation in genome size is associated with the depth distribution of zooxanthellate corals, as has been observed in reef fishes (15). The Caribbean genus *Agaricia* is an interesting subject to investigate this relationship because it consists of species that are found at different depth ranges with some considerable overlaps (30).

One major difficulty in conducting comparative studies among scleractinian corals is that their taxonomy is presently in a state of flux, with molecular studies revealing many discrepancies with morphology-based species delimitations (31). In *Agaricia*, high levels of incongruence between morphology and *atp6* mitochondrial DNA sequences were recorded (30). Other mitochondrial markers (*nad5* and *cox1*-1-rRNA) have also been unable to differentiate closely-related *Agaricia* species (32) as can be expected for mitochondrial genes which are known to evolve too slowly for species delimitation in most anthozoans (33, 34). Thus, there is a need to use nuclear markers in defining species boundaries within *Agaricia*.

This study mainly aimed to address the following research questions:

- Do genome sizes vary significantly between *Agaricia* coral species?
- Are variations in genome size within and among species correlated with genetic polymorphism (*π*)?
- Are variations in genome sizes correlated with depth?

Considering their distinct depth distributions, we would expect genome sizes to vary among *Agaricia* species and allow us to discriminate among them. Here, we try to account for the possible influence of effective population size (Ne) using nucleotide polymorphism (*π*) as a proxy (10, 35). We would expect genome sizes to be negatively correlated with *π* (10) and positively correlated with collection depth due to the relaxed selection that could be associated with the stability of deep-water environments (28, 29). To properly interpret this variation in genome size in a taxonomic context, we first attempt to genetically delimit the *Agaricia* samples by sequencing the nuclear markers internal transcribed spacer 2 (ITS2) and L-threonine 3-dehydrogenase (TDH) and analyzing the sequences with different approaches to species delimitation (36).

Understanding the variation in genome size within and between species of the Caribbean coral genus *Agaricia* will provide insights into the evolution of genome size in the marine environment and how depth-related environmental factors are linked to this biological trait. Moreover, this study provides additional genome size data necessary for a wider phylogenetic comparative analysis within reef-building scleractinian corals.

## Materials and Methods

### Study area and sample collection

Curaçao is an island in the southern Caribbean surrounded by fringing reefs around 20 m to 250 m away from the coast. The reefs located on the leeward side of the island are generally characterized by a 50-200 m wide reef flat with patchy coral communities followed by a steep drop-off at around 7-15 m depth where coral cover and diversity are the highest (37). In this study, the two collection sites Director’s Bay (12°3.9585’ N, 68°51.5955’ W) and Snakebay (12°8.3496’ N, 68°59.83956’ W) are both located on the leeward side of the island (Figure 1). These two sites on the same side of the island were selected as they are expected to harbor similar coral species compositions that would allow the collection of the same species across the depth range. Reefs on the windward side of the island are generally poorly developed and mostly dominated by *Sargassum* due to strong wave action (24).

**Fig. 1.**
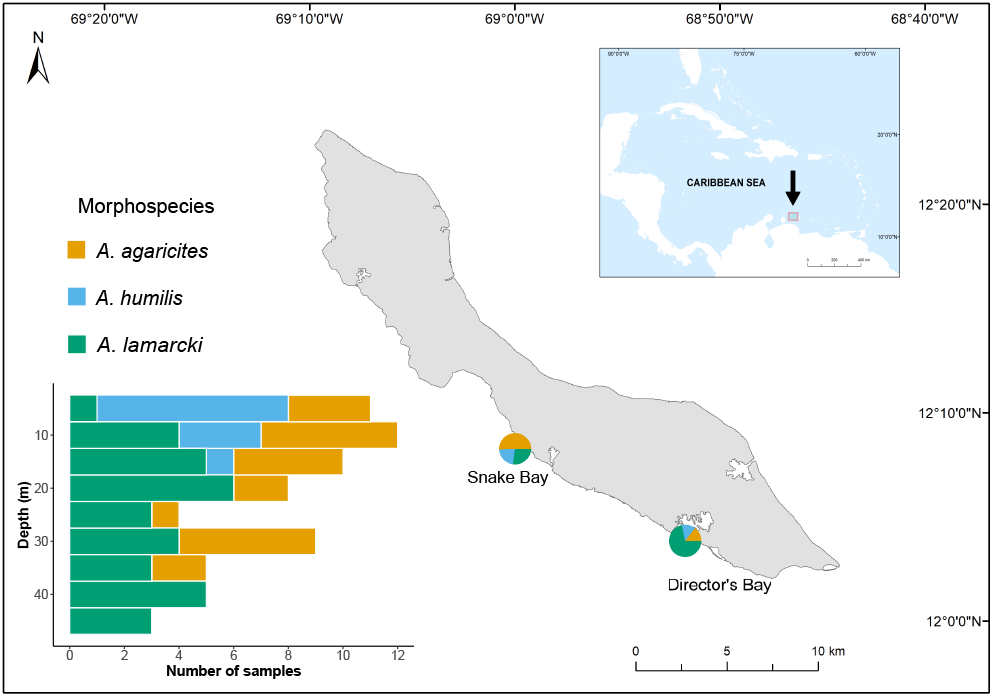
Map of the study area: Curaçao in southern Caribbean. Pie charts correspond to the proportion of samples for the three morphospecies collected from the two sites. The depth distribution profile of the collected samples from the two sites combined are shown on the bottom left. Number of samples collected per species: *A. lamarcki*: n = 34; *A. agaricites*: n = 22; *A. humilis*: n = 11. The difference in the relative frequencies of each morphotype collected between the two sites could be attributed to differences in the topography of the reef sites and the maximum sampling depth. The map is generated with the ArcGIS software (Redlands, 2011), using shapefiles obtained from www.geominds.de

*Agaricia* samples were collected from the two sites in December 2021 using scuba diving from 47 m depth (in Director’s Bay) or 35 m depth (in Snake Bay) up to the surface. Different morphotypes present along the depth gradient were sampled randomly. A total of 67 samples were collected at these sites (Director’s Bay: n = 35; Snake Bay: n = 32): each colony was photographed and its depth recorded, then a small fragment (*∼* 2 cm^2^) was broken off. The specimens were brought back to the Caribbean Research and Management of Biodiversity (CARMABI) research station and preserved in 96% ethanol. They were then transported to the Evolutionary Biology and Ecology (EBE) research unit at the Université libre de Bruxelles (ULB) and stored at -20°C for further processing.

### Morphospecies identification

Of the seven *Agaricia* morphospecies found in the Caribbean, five are reported to occur on Curaçao reefs. Two morphospecies are considered depth generalists (*A. lamarcki* and *A. agaricites*), one is a shallow-water (<30 m) specialist (*A. humilis*), and two are mesophotic (30-150 m) specialists (*A. grahamae* and *A. undata*). Morphospecies were identified based on the skeletal morphology of the collected fragment following Veron et al. (2022) (38) and Bongaerts et al. (2013) (30), aided by the live underwater photographs of the colonies (Figure 6).

### Genetic analyses

Genomic DNA was extracted from the tissue samples following the protocol of Macherey-Nagel’s Nucleospin Tissue Kit. The quantity and purity of the DNA extracts were assessed with a Nanodrop spectrophotometer. Amplification of ITS2 and TDH nuclear markers was performed in 25 µL PCR reaction mixes containing 12.5 µL DreamTaq Green PCR Master Mix, 10 µL ultrapure water, 1 µL reverse and 1 µL forward primer, and 0.5 µL DNA. Coral-specific ITS2 primers (39) used were : ITSc2-5 5’-AGCCAGCTGCGATAAGTAGTG–3’ (forward primer) and R28S1 5’-GCTGCAATCCCAAACAACCC-3’ (reverse primer). The TDH primers that were designed using the available transcriptome of *A. lamarcki* (40) were: 5’-CACAATCCAGAGACCAAAAACA-3’ (forward primer) and 5’-TCCAAGCCAAACTTGTGATG-3’ (reverse primer). PCR conditions for both ITS2 and TDH markers consisted of an initial denaturation at 95°C, followed by 45 cycles of 30 s denaturation at 95°C, 30 s annealing at 53°C, and 60 s elongation at 72°C. Successful PCR products were sent to Macrogen in Amsterdam for bidirectional Sanger sequencing.

Forward and reverse sequences were assembled and cleaned using Sequencher v5.4.6 (GeneCodes, USA). Length-variant heterozygotes were phased using Champuru (41). As two individuals (DMR-064 and DMR-015) could not be phased, we resorted to single-molecule Nanopore sequencing to resolve the haplotypes of one of them (DMR-064), which allowed us to deduce the most likely phasing for the other one (DMR-015). Briefly, the amplicon was spiked into one LSK114 library that was run on a Flongle R10.4.1 flow cell. After base-calling using guppy 6.3.8, matching reads were recovered by aligning them using minimap2 (42) against the known sequence of the marker, imported into Sequencher, and scutinized at the two SNP positions to deduce the haplotypes of the individual. All phased haplotypes were aligned using the E-INS-i method in the online implementation of MAFFT v7 (43).

A haploweb of the ITS2 and TDH sequences was constructed using HaplowebMaker (44) with default parameters, treating indels as a 5th character and ambiguous characters as errors. This method relies on the co-occurrence of alleles in a group of individuals to delimit genetic pools corresponding to putative species based on a single molecular marker (45).

A distance-based approach to species delimitation was carried out with the Assemble Species by Automatic Partitioning (ASAP) web tool, using JC69 Jukes-Cantor as a substitution model (46). A maximum likelihood phylogenetic tree was also constructed using the online implementation of IQ-TREE (47), with JC+I as the best-suited substitution model for ITS2 and HKY+F+G4 for TDH (48).

Nucleotide polymorphism *π* for each species was calculated using the nuc.div function from pegas (49) package in R.

### Genome size estimation

Genome sizes were measured using the improved Feulgen protocol of M.Tawfeeq et al. (in prep.). In brief, the protocol includes fixation of the tissues on the microscopic slide, hydrolysis with acid, staining (with Schiff’s reagent), and rinsing of the slides to remove any unbound stain (50). Three standards with known C-values were used for each Feulgen experiment: *Lasius niger* (0.31 pg) (JF Flot, unpublished data), *Periplaneta americana* (3.41 pg) (51), and *Allium schoenoprasum* (7.7 pg) (52). A ToupCam camera mounted on a Leitz Laborlux D microscope was used to view the slides. Nuclei measurements were performed using ImageJ, an open-source program developed by the National Institute of Health (NIH). Three important measurements were obtained: the area of the nucleus, the mean grey value of the nucleus, and the mean grey value of the background. These values are used to calculate the optical density (OD), calculated as the difference between the mean grey value of the background and the mean gray value of the nucleus. This OD value is multiplied per area of the nucleus to calculate the integrated optical density (IOD), which is used for the calculation of the C-value (amount of DNA per haploid cell) for each specimen against the three standards with known C-value. For each sample, at least 30 nuclei of different sizes and compaction levels were measured for a reliable estimation. The coral samples consisted mostly of smaller round nuclei that tended to be more compact and common, whereas bigger and less compact ones occurred at low densities. Elongated nuclei were also present. These three nuclei types gave comparable IODs. The nuclei of the algal symbiont were distinguished from those of the coral by visually inspecting the presence of a slightly stained cell wall around the latter (53).

### Statistical analyses

Parametric assumptions were evaluated with Levene’s test for homogeneity of variances (54) and Shapiro-Wilk to test for normality (55). After all parametric assumptions were met, a one-way ANOVA was conducted to test whether mean genome size differed among species using the aov function in stats package (56). To identify which species are statistically different, a posthoc Tukey test (57) was performed. Pearson’s correlation analyses were conducted to test the relationships between nucleotide polymorphism (*π*) and the level of intraspecific variation in genome size (represented by the variance); nucleotide polymorphism (*π*) and average genome size per species; average depth and average genome size; and collection depth and genome size estimates of depth-generalists species (*A. lamarcki* and *A. agaricites*). Parametric assumptions (linearity and normality) were first tested before conducting the analyses. All statistical analyses were done in R (56) using the Rstudio console (58).

## Results

In the present study, only three morphospecies (*A. lamarcki, A. agaricites* and *A. humilis*) were identified. The depth distribution of the collected samples (from two sites combined) and their proportions in each site are shown in Figure 1.

### Species delimitation

All 67 coral samples were successfully sequenced for the nuclear ITS2 and TDH markers. For both markers, 3 allele pools corresponding to the three morphospecies identified were observed in the haploweb analyses. The largest allele pool corresponds to *A. lamarcki*, followed by *A. agaricites* and *A. humilis* (Figure 2 A,D). ASAP analysis of ITS2 sequences supported the best partition consisting of 3 species (Figure 2C) (*A. lamarcki, A. agaricites*, and *A. humilis*, whereas the best partition for the TDH sequences revealed 2 species (Figure 2F), grouping *A. agaricites* and *Agaricia humilis* together. The maximum likelihood tree for both markers also supports three species among the collected samples (Figure 2 B,E).

**Fig. 2.**
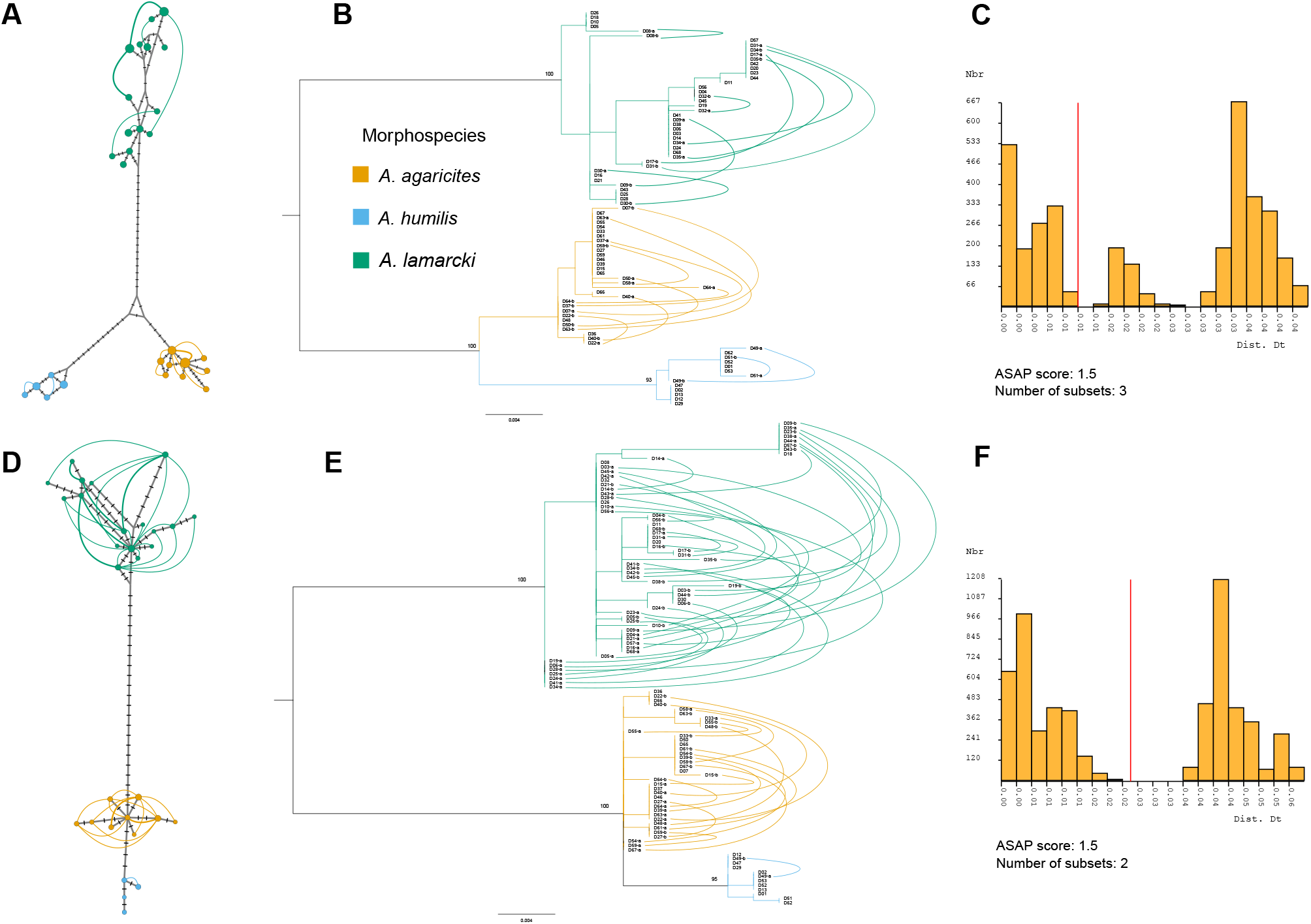
Different approaches to molecular species delimitation of Agaricia samples using the ribosomal ITS2 marker (A-C) and nuclear TDH marker (D-F). **(A)** Haploweb of ITS2 sequences colored according to morphospecies (DMR-005 and DMR-010, the two individuals initially attributed putatively to *A. grahamae*, turned out to be deep-water morphs of *A. lamarcki* and are therefore shown in green). The circles represent the haplotypes with the diameter proportional to the number of individuals harboring each of them, while curves connect co-occurring haplotypes in a heterozygous individual (width of the curve is proportional to the number of heterozygotes harboring the two haplotypes). **(B)** Midpoint-rooted ITS2 gene tree. Bezier curves connect haplotypes co-occuring in heterozygous individuals. Values on the branches represent bootstrap values. **(C)** Histogram of the 3-species partition best supported by ASAP. **(D)** Haploweb of TDH sequences showing 3 allele pools, colored according to morphospecies. **(E)** Midpoint-rooted TDH gene tree with curves connecting co-occuring haplotypes. **(F)** Histogram of the best (2 species) partition for the TDH sequences suggested by ASAP.

We observed two colonies whose appearance did not clearly conform with the morphospecies descriptions and identified them tentatively as *A. grahamae* (a mesophotic specialist) based on their collection depths. With their sloping ridges, the colonies look most similar to *A. lamarcki* than to any other *Agaricia* species. However, their striking septo-costae, less obvious white-colored mouths, and their coloration make them quite different from the “normal” *A. lamarcki* morphotypes (Supplementary Feigure 6E, F). These two samples were grouped together with the rest of the *A. lamarcki* samples based on their ITS2 and TDH sequences. For this reason, we treated them as deep-water morphs of *A. lamarcki*.

Figure 5B shows no significant differences in the haplotype composition between the two collection sites since frequent haplotypes are shared between them. Thus, samples from the two sites were pooled together for the rest of this study.

### Genome size measurements

The genome sizes of *Agaricia* specimens analyzed here range from 0.359 to 0.593 pg with the largest mean genome size recorded in *A. agaricites* (mean ± SD: 0.495 pg ± 0.034 pg), followed by *A. lamarcki* (0.448 ± 0.047 pg), and then by *A. humilis* (0.434 pg ± 0.036). The mean genome size of these three morphospecies were significantly different (ANOVA df = 2, F = 11.29, p < 0.05) (Figure 3C). The posthoc Tukey test revealed that the mean genome size of *A. agaricites* differed significantly from that of *A. lamarcki* (p-adjusted < 0.05) and *A. humilis* (p-adjusted < 0.05). On the other hand, there was no significant difference in genome size between *A. lamarcki* and *A. humilis* (p-adjusted = 0.611).

**Fig. 3.**
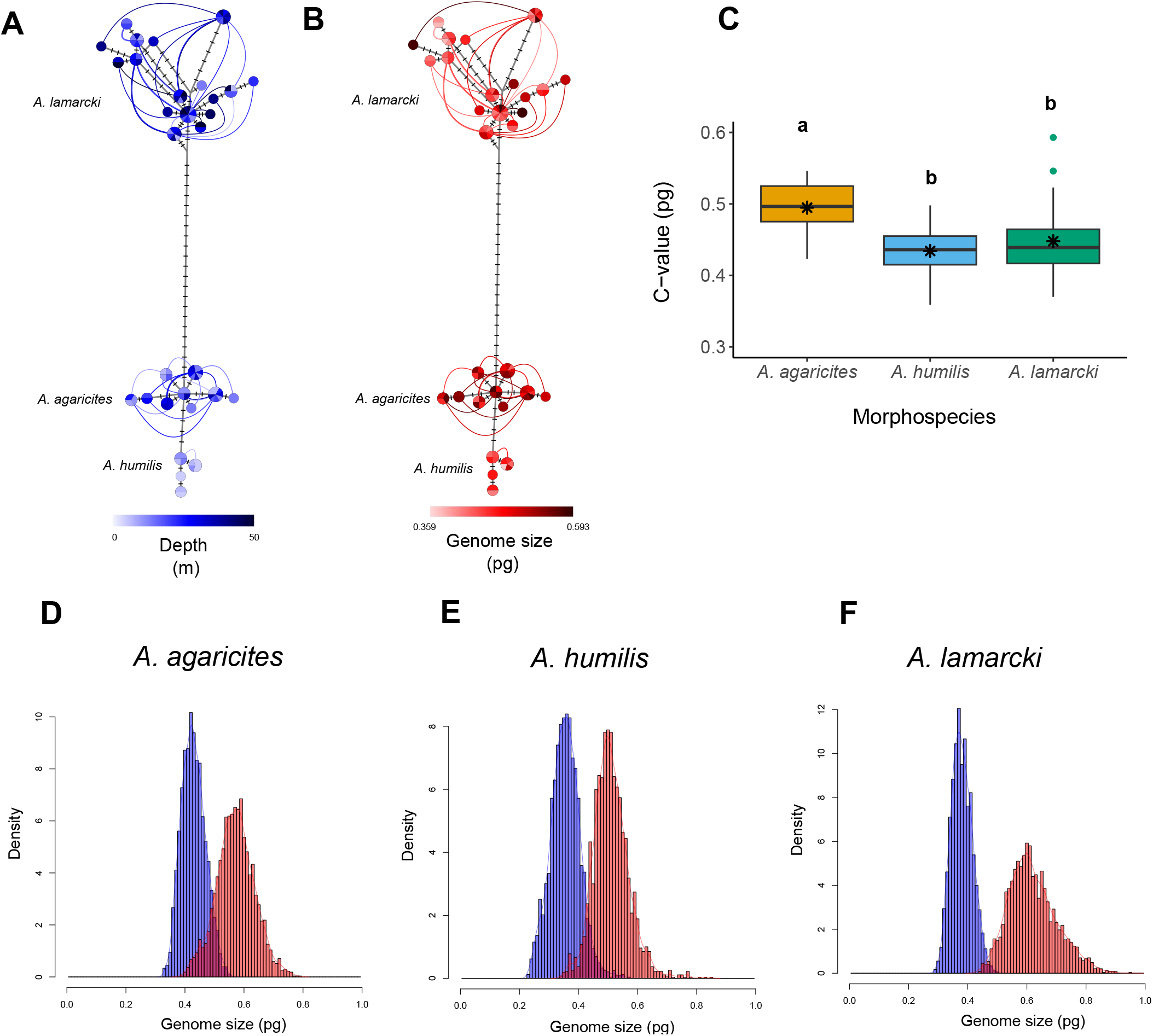
Intra- and interspecific genome size variation in *Agaricia* samples analyzed in this study. **(A-B)** Haplowebs of TDH sequences colored according to **(A)** depth in meters and **(B)** genome size (pg) estimated using FIAD. **(C)** Interspecific genome size variation (asterisk represents mean genome size, different lowercase letters represent statistical significance with α = 0.05). **(D-F)** Intraspecific genome size variation represented by histograms of the minimum (blue) and maximum (red) genome size values for each species.

*A. agaricites* exhibits the smallest range of genome size (0.423 to 0.546; 1.3-fold variation), followed by *A. humilis* (0.359 to 0.498 pg; 1.4-fold variation) and *A. lamarcki* (0.396 to 0.593 pg; 1.5-fold variation) (Figure 3D-F).

For the ITS2 marker, nucleotide polymorphism (*π*) was highest in *A. lamarcki* (*π* = 0.00535), followed by *A. humilis* (*π* = 0.00311), and lowest in *A. agaricites* (*π* = 0.002209). A non-significant negative correlation (Pearson’s r = -0.555, p = 0.625) was recorded between *π* and mean genome size (Figure 4A). A clear positive correlation (Pearson’s r = 0.984) between *π* and levels of intraspecific variability in genome size was recorded (Figure 4B), however, this relationship was not significant (p = 0.113), with alpha = 0.05. Non-significant positive correlations between nucleotide polymorphism and genome size mean (Pearson’s r = 0.117, p = 0.926) and variance (Pearson’s r = 0.863, p = 0.337) were recorded for the TDH marker (Figure 4C,D).

**Fig. 4.**
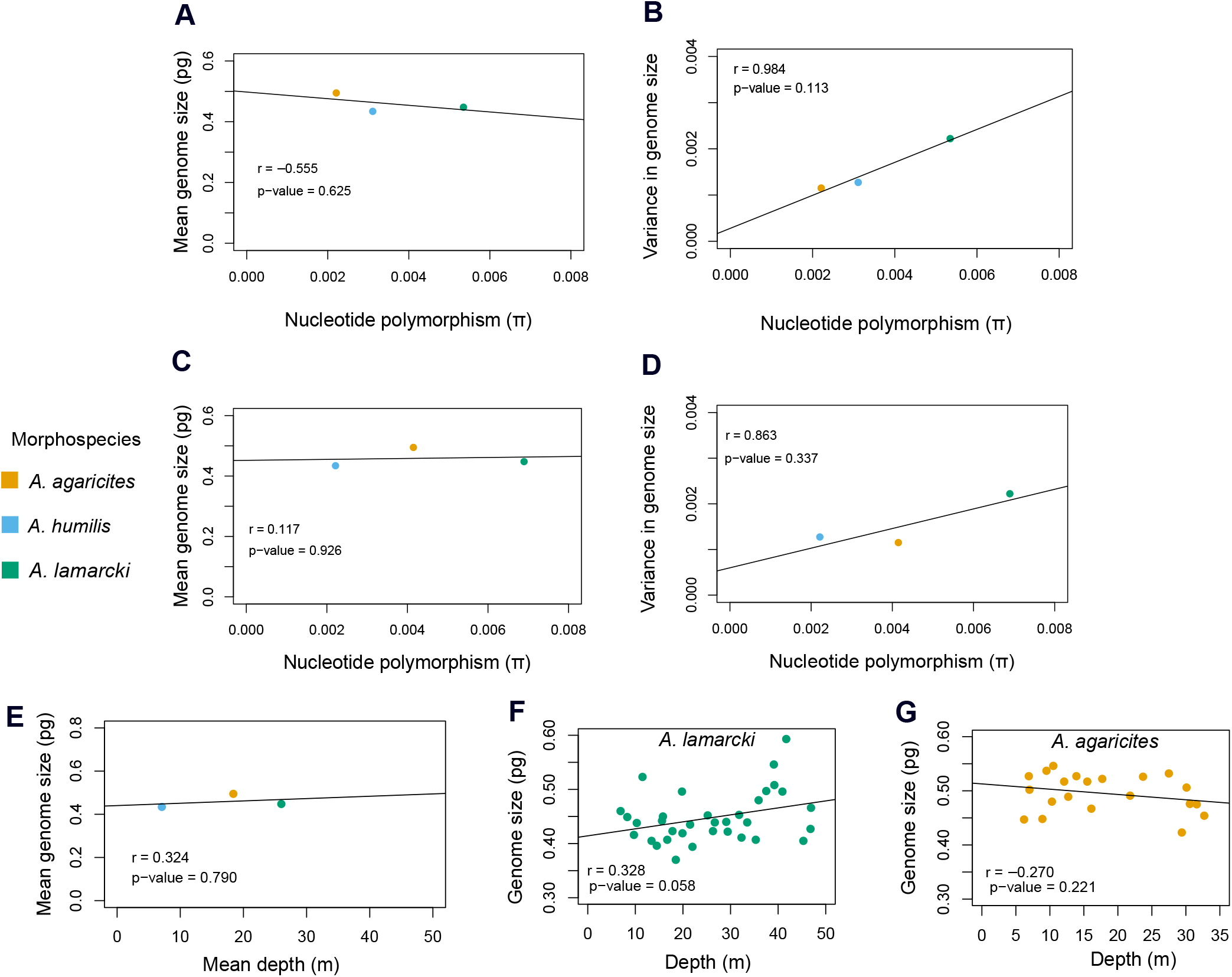
Relationship between nucleotide polymorphism π and mean genome size (pg), and intraspecific variation in genome size (represented by variance) for **(A-B)** ITS2 and **(C-D)** TDH marker. Scatterplots between: **(E)** mean genome size (pg) and mean depth (m); genome size (pg) and collection depth (m) for the depth-generalists **(F)** *A. lamarcki* and **(G)** *A. agaricites*. r refers to Pearson’s correlation coefficient, α = 0.05r refers to Pearson’s correlation coefficient, α = 0.05

**Fig. 5.**
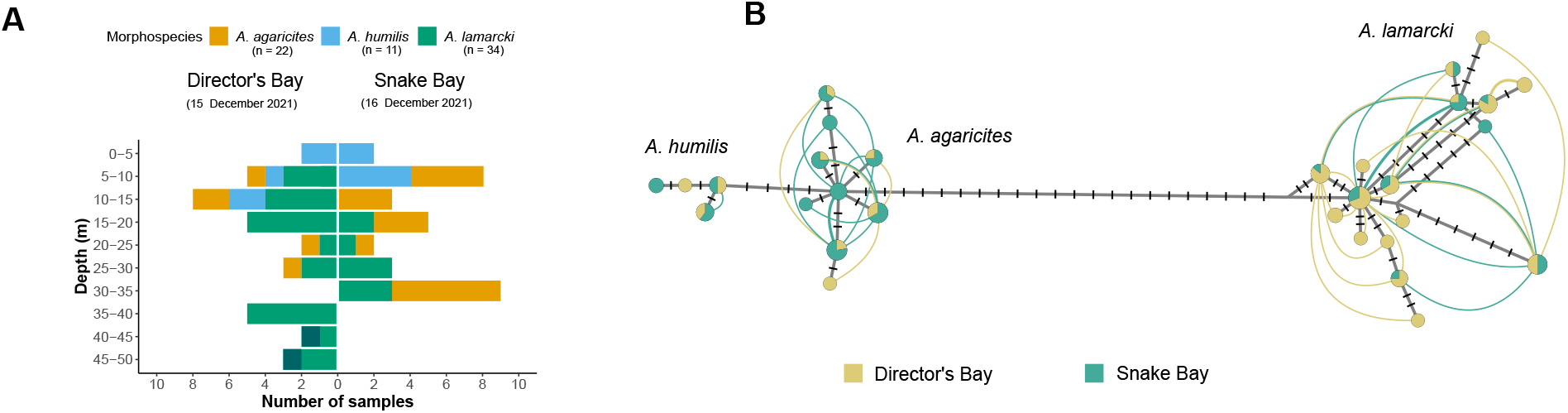
*Agaricia* morphospecies collected from the two sampling sites (Director’s Bay and Snake Bay). (**A**) Proportion of samples collected along the depth range in both sites (**B**) Haploweb of TDH sequences colored according to the collection site, showing genetic connectivity between the two sites.

**Fig. 6.**
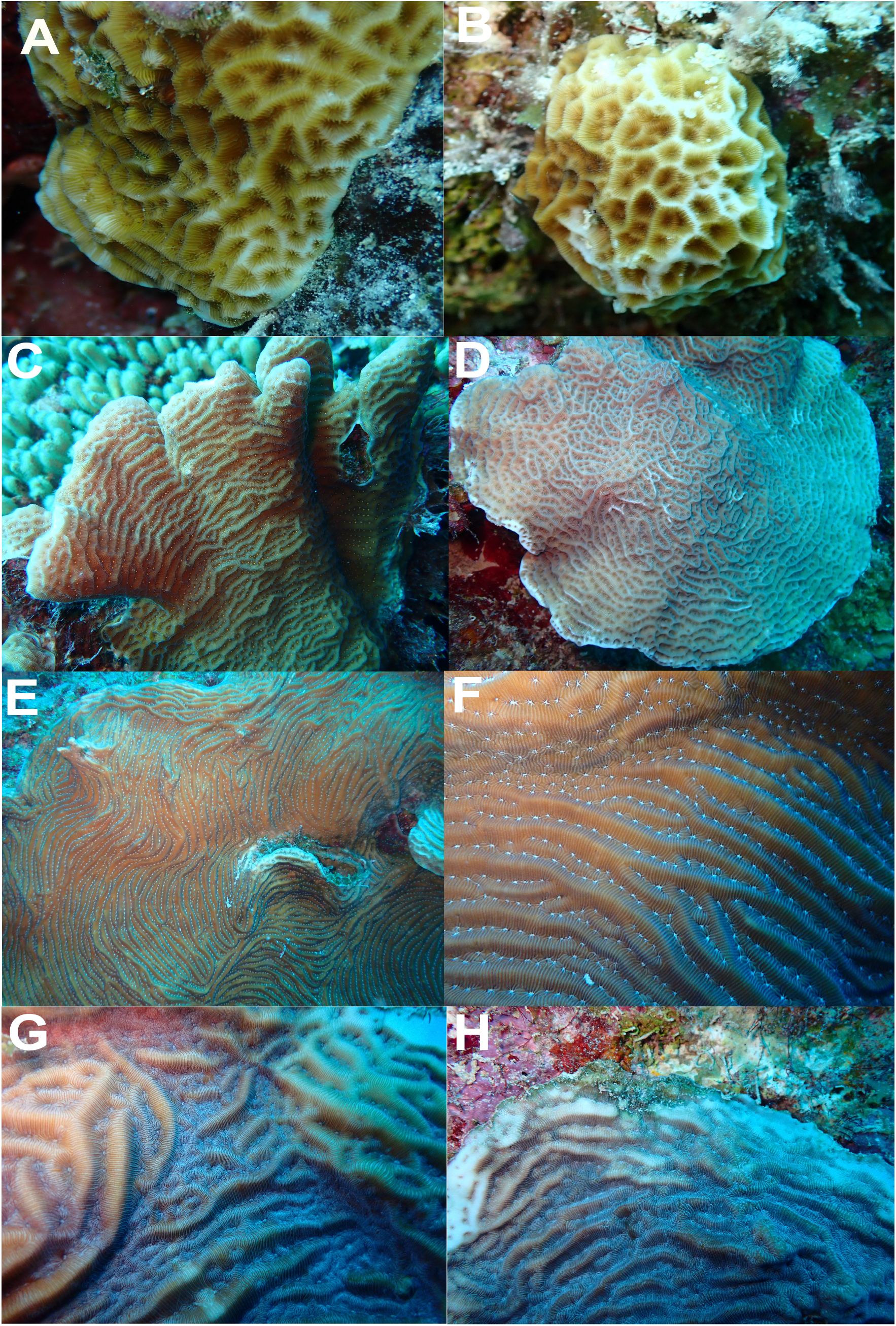
*Agaricia* morphospecies collected from Curaçao. (**A-B**) *A. humilis*, (**C**) *A. agaricites* forming fronds, (**D**) *A. agaricites* with plating morphology, (**E-F**) *A. lamarcki*, (**G-H**) deep-water morphs of *A. lamarcki*

## Discussion

### Morphology and two nuclear markers concur in delimiting three *Agaricia* species among our samples

Previous studies using mitochondrial markers have been unsuccessful in delineating the species boundaries in this coral genus. For example, the mitochondrial *atp6* marker has only been able to distinguish two major lineages in *Agaricia*: shallow-water (*A. agaricites* and *A. humilis*) and deep-water (*A. lamarcki* and *A. grahamae*) lineages, but with some *A. humilis* individuals grouping with *A. lamarcki* and *A. grahamae*. Morphological species boundaries, therefore, were incongruent with this genetic marker. The nuclear 28S sequences support the division of this genus into two lineages and further differentiate between *A. agaricites* and *A. humilis* (shallow species) and *A. lamarcki* and *A. grahamae* (deeper species) (30).

In this study, we found that the nuclear markers ITS2 and TDH and morphology are congruent in delimiting 3 *Agaricia* species among the samples collected: one shallow-water specialist (*A. humilis*), one depth generalist (*A. lamarcki*), and one with a depth distribution intermediary between the two (*A. agaricites*). The results suggest that these 3 *Agaricia* taxa are good biological species with no evidence of gene flow, as supported by the different species delimitation analyses conducted.

The faster coalescence rate of ITS2 due to its concerted evolution (59, 60), its relatively high mutation rate and the availability of a primer pair (61) that works on all coral species so far make it a good marker of choice for single-locus delimitation in corals. In *Stylophora*, ITS2 has allowed to find out which morphotypes were simply deep-water morphs of common species and which ones were distinct species (62). It has also unveiled the real species boundaries that were obscured by phenotypic plasticity in *Pocillopora* (39). In this study, ITS2 has also revealed 3 species that are congruent with morphological descriptions. The TDH marker has been recently used by Ramírez-Portilla et al. (2022) to delimit corals in the genus *Acropora* (63). This study suggests that TDH could also be a good marker of choice for species delimitation in *Agaricia*. Several young *Agaricia* colonies sampled in this study cannot be confidently identified based on the available morphospecies descriptions in the literature. For example, some samples that are genetically grouped with *A. humilis* for both markers may be morphologically identified as young colonies of *A. agaricites*. In the future, providing detailed information on the morphological variability of different species delineated using molecular data would be helpful in identifying *Agaricia* morphospecies.

### Genome size measurements reveal unexpected intra-as well as interspecific variations among *Agaricia* corals in Curaçao

Genome sizes of 3 *Agaricia* species estimated using FIAD showed a 1.7-fold variation within this genus, with the smallest genome size recorded in the shallow-water specialist *A. humilis* (0.359 pg) and the largest one in the deep-water morph of *A. lamarcki* (0.593 pg). The highest average genome size (0.495 pg) and lowest intraspecific variation (1.3-fold) were recorded in the depth generalist *A. agaricites*. The average genome size of *A. lamarcki* (0.448 pg) and *A. humilis* (0.434 pg) did not differ significantly.

This study provides the first exploration of intraspecific genome size variation in scleractinian corals and contributes to the few studies that reported this microevolutionary aspect of genome size variation. In rotifers, the twofold variation in genome size within the same population was attributed to copy number variable regions (CNVs) which are mainly composed of satellite DNA repeats (21). Different genome sizes were also observed between sexes in orthopteran insects, with females having larger genomes due to the bigger X chromosome (20, 22). The intraspecific variation observed here is comparable to that reported in fungus-forming ants (64) and snapping shrimps (19), where significant genome size variation among geographic locations and even within the same colony (in snapping shrimps) was also recorded. This variation has been suggested to be influenced by differences in the proliferation and deletion of transposable elements among populations (19). In fungus-forming ants, the variation in genome size among populations was attributed to such modifications which are suggested to be triggered by stressful conditions during the dispersal of species to new habitats (64). To understand the specific mechanisms and identify genomic elements associated with the differences in genome size of the *Agaricia* species studied here, individuals with extreme genome sizes from each species should be selected and sent for whole genome sequencing.

The positive association between average genome size and niche breadth reported by dos Santos et al. (15) leads to the expectation that depth-generalist species would in general, have larger genome size than a species restricted to a narrower depth range. Here, although the average genome size of the depth-generalist *A. lamarcki* (0.448 pg) is a bit larger than the depth-specialist *A. humilis* (0.434 pg), this difference is not statistically significant. The largest genome size was recorded in another depth-generalist species (*A. agaricites*) with an intermediate depth distribution. Moreover, the highest intraspecific variation in genome size was recorded in *A. lamarcki* (1.5-fold variation), the species with the widest depth range, which suggest that higher intraspecific variability in genome size might be associated with wider ecological niches.

While variation in genome size in eukaryotes has been mainly attributed to the amount of non-coding DNA (e.g. transposable elements) (65, 66), gene expansions may also contribute to this variation and could be a source of novel gene functions (67). In *Acropora*, analysis of whole genome sequences of 15 species has shown gene expansions in their common ancestor which might have contributed to the adaptations of this genus during past warming events and facilitated their wide distributions (68). Gene family expansions related to immune functions have also been reported in *Stylophora pistillata, Acropora digitifera, Orbicella faveolata*, and *Pocillopora damicornis* (69, 70). Moreover, a recent study has confirmed whole genome duplications of the coral symbiont *Durusdinium trenchii*, which is known to confer thermal tolerance to the host coral (71). These studies suggest that genome size evolution (of the coral host or algal symbiont) may play a key role in the adaptive potential of corals and that whole genome assemblies of a wider range of taxa with different environmental tolerances could provide a better understanding of this relationship.

Nucleotide polymorphism (*π*), as a proxy for Ne (10), is positively correlated with the levels of intraspecific variation in genome size, although this relationship is not significant. The relatively high p-value reported in the analysis is probably due to the low number of species included in the study. This could be addressed in the future by including other *Agaricia* species and other genera to see whether this trend holds. Thus, the hypothesis of reduced polymorphism (and thus reduced effective population size) in species with larger genomes cannot be confidently supported by the results of the current study.

### No correlation between depth and genome size was detected

This study showed non-significant and weak positive (*A. lamarcki*) (Pearson’s r = 0.328, p = 0.058) (Pearson’s r = -0.272, p = 0.221) (Figure 4F) and negative (*A. agaricites*) (Pearson’s r = -0.272, p = 0.221) (Figure 4G) correlations between genome size and collection depth in the two depth-generalist species. Moreover, interspecific differences in genome size also do not support the hypothesis that the average genome size of coral species is positively linked to its average depth (Pearson’s r = 0.109, p = 0.382) (Figure 4E). Thus, the hypothesis of increased genome size with depth as a result of less efficient selection cannot be confidently supported in this study due to this very weak and non-significant relationship. The inclusion of the mesophotic specialist *A. grahamae* in this analysis might shed light as to what extent the genome size of a species varies with contrasting depth-related ecological factors. It will also be interesting to test this hypothesis at other locations using other coral genera.

The few studies that reported intraspecific variation in genome size (19, 21, 64) did not investigate whether this variation is linked to any ecological patterns. The level of intraspecific variability observed here seems to be slightly in accordance with the variability in depth. For example, *A. lamarcki*, the most generalist in terms of depth, also has the highest level of intraspecific variability in genome size which could have potential implications regarding the range of depth-related ecological factors it can tolerate (i.e. ecological flexibility).

In marine fishes, a significant positive correlation between genome size and maximum depth was recovered in only one subclade (Labriformes) while non-significant correlations between these two variables are common, especially after correcting for phylogenetic non-independence (72). This “modest” evidence of genome size increase with depth reported in marine fishes was corroborated by the same pattern in crustaceans, which they attributed to the passive accumulation of redundant genomic elements in the environmentally stable deep-water habitats (72–74).

## Conclusions and perspectives

In summary, we found that the genome size of 3 *Agaricia* species partly varies, significantly discriminating only one species (*A. agaricites*) with an intermediate depth distribution. Nucleotide polymorphism (as a proxy for Ne) was not significantly correlated with average genome size and intraspecific variability There is also no evidence of a positive correlation between intraspecific genome sizes and their collection depths for both depth generalists *A. lamarcki* and *A. agaricites*, as well as the average genome size of each species and their average collection depth. In addition to our main findings, our study also revealed concordant patterns between morphological species descriptions and nuclear ITS2 and TDH sequences in delimiting three *Agaricia* species. Establishing species boundaries is important to know whether two morphotypes are just morphs or different species. Here, the different species delimitation approaches strongly suggest that the two individuals initially suspected of belonging to *A. grahamae* may just be growth forms of *A. lamarcki*.

As genome size data become available for more coral species, future phylogenetic comparative studies that take into account a wider taxonomic spectrum might reveal a tighter link between the relationships investigated here. This study contributes to what is known about the extent of genome size variability within a species and suggests that this variation might be more common than what is reported, and therefore warrants more attention. The increasing availability of reference whole genome sequences even for non-model organisms could improve our knowledge about the mechanisms underlying genome size variation within and between species.

## ACKNOWLEDGEMENTS

We thank Astrid Verkek and Jeroen Schneider for their assistance during sample collection and Alice Salussolia for her help in the laboratory. This study was supported by Fonds de la Recherche Scientifique—FNRS (research grant J.0272.17 to Jean-François Flot).

## Supplementary Material

**Table 1.** Genome sizes of scleractinian corals estimated using flow cytometry (FCM), Feulgen image analysis densitomety (FIAD), and/or next generation sequencing (NGS). Note: Genome size data were standardized by converting assembly size (Mb) to picograms (pg), where 1pg = 978 Mb (75).

| Species | C-value (pg) | Assembly size (Mb) | Method | Reference |
| --- | --- | --- | --- | --- |
| *Acropora digitifera* | 0.43 |  | FCM | Shinzato et al. (2021) (68) |
| *Catalaphyllia jardinei* | 0.9 |  | FCM | Adachi et al. (2017) (8) |
| *Euphyllia ancora* | 0.64 |  | FCM | Adachi et al. (2017) (8) |
| *Euphyllia divisa* | 0.72 |  | FCM | Adachi et al. (2017) (8) |
| *Physogyra lichtensteini* | 0.98 |  | FCM | Adachi et al. (2017) (8) |
| *Siderastrea stellata* | 1.14 |  | FIAD | Gregory (2022) (18) |
| *Acropora acuminata* | 0.40 | 395 | NGS | Shinzato et al. (2021) (68) |
| *Acropora awi* | 0.44 | 429 | NGS | Shinzato et al. (2021) (68) |
| *Acropora cytherea* | 0.44 | 426 | NGS | Shinzato et al. (2021) (68) |
| *Acropora digitifera* | 0.43 | 416 | NGS | Shinzato et al. (2021) (68) |
| *Acropora echinata* | 0.41 | 401 | NGS | Shinzato et al. (2021) (68) |
| *Acropora florida* | 0.45 | 442 | NGS | Shinzato et al. (2021) (68) |
| *Acropora hyacinthus* | 0.46 | 447 | NGS | Shinzato et al. (2021) (68) |
| *Acropora intermedia* | 0.43 | 417 | NGS | Shinzato et al. (2021) (68) |
| *Acropora microphthalmia* | 0.39 | 384 | NGS | Shinzato et al. (2021) (68) |
| *Acropora muricata* | 0.43 | 421 | NGS | Shinzato et al. (2021) (68) |
| *Acropora nasuta* | 0.43 | 416 | NGS | Shinzato et al. (2021) (68) |
| *Acropora selago* | 0.40 | 393 | NGS | Shinzato et al. (2021) (68) |
| *Acropora tenuis* | 0.41 | 403 | NGS | Shinzato et al. (2021) (68) |
| *Acropora yongei* | 0.45 | 438 | NGS | Shinzato et al. (2021) (68) |
| *Acropora millepora* | 0.40 | 387 | NGS | Ying et al. (2018) (76) |
| *Acropora gemmifera* | 0.41 | 401 | NGS | Shinzato et al. (2021) (68) |
| *Astreopora myriophthalma* | 0.38 | 373 | NGS | Shinzato et al. (2021) (68) |
| *Montipora efflorescens* | 0.66 | 643 | NGS | Shinzato et al. (2021) (68) |
| *Montipora cactus* | 0.67 | 653 | NGS | Shinzato et al. (2021) (68) |
| *Montipora capitata* | 0.63 | 614 | NGS | Helmkamp et al. (2019) (77) |
| *Stylophora pistillata* | 0.41 | 400 | NGS | Voolstra et al. (2017) (69) |
| *Orbicella faveolata* | 0.50 | 486 | NGS | Prada et al. (2016) (78) |
| *Pocillopora damicornis* | 0.24 | 234 | NGS | Cunning et al. (2018) (70) |
| *Pocillopora verrucosa* | 0.39 | 380 | NGS | Buitrago-López et al. (2020) (79) |

**Table 2.** Genome size of *Agaricia* samples estimated using the refined Feulgen Image Analysis Densitometry (FIAD) method (M.Tawfeeq, in prep) (*individuals we considered as “deep-water morphs” of *A. lamarcki*).

| Lab ID | Morphospecies | Collection site | Depth (m) | Genome size (pg) | CV |
| --- | --- | --- | --- | --- | --- |
| DMR-007 | *A. agaricites* | Director's Bay | 7 | 0.502 | 15% |
| DMR-015 | *A. agaricites* | Director's Bay | 12.7 | 0.489 | 6% |
| DMR-022 | *A. agaricites* | Director's Bay | 29.4 | 0.423 | 9% |
| DMR-027 | *A. agaricites* | Director's Bay | 10.3 | 0.480 | 9% |
| DMR-033 | *A. agaricites* | Director's Bay | 21.8 | 0.491 | 10% |
| DMR-036 | *A. agaricites* | Snake Bay | 32.7 | 0.454 | 8% |
| DMR-037 | *A. agaricites* | Snake Bay | 31.6 | 0.475 | 10% |
| DMR-039 | *A. agaricites* | Snake Bay | 30.6 | 0.476 | 10% |
| DMR-040 | *A. agaricites* | Snake Bay | 30.1 | 0.506 | 8% |
| DMR-046 | *A. agaricites* | Snake Bay | 13.9 | 0.527 | 6% |
| DMR-048 | *A. agaricites* | Snake Bay | 8.9 | 0.448 | 6% |
| DMR-050 | *A. agaricites* | Snake Bay | 6.2 | 0.447 | 10% |
| DMR-054 | *A. agaricites* | Snake Bay | 10.5 | 0.546 | 10% |
| DMR-055 | *A. agaricites* | Snake Bay | 9.5 | 0.537 | 5% |
| DMR-058 | *A. agaricites* | Snake Bay | 17.7 | 0.522 | 5% |
| DMR-059 | *A. agaricites* | Snake Bay | 15.5 | 0.517 | 6% |
| DMR-061 | *A. agaricites* | Snake Bay | 6.9 | 0.527 | 6% |
| DMR-063 | *A. agaricites* | Snake Bay | 23.7 | 0.526 | 10% |
| DMR-064 | *A. agaricites* | Snake Bay | 12.1 | 0.523 | 9% |
| DMR-065 | *A. agaricites* | Snake Bay | 16.1 | 0.467 | 11% |
| DMR-066 | *A. agaricites* | Snake Bay | 27.5 | 0.532 | 9% |
| DMR-067 | *A. agaricites* | Snake Bay | 30.6 | 0.476 | 12% |
| DMR-001 | *A. humilis* | Director's Bay | 4.5 | 0.456 | 10% |
| DMR-002 | *A. humilis* | Director's Bay | 4.9 | 0.446 | 9% |
| DMR-012 | *A. humilis* | Director's Bay | 9.8 | 0.414 | 12% |
| DMR-013 | *A. humilis* | Director's Bay | 10.4 | 0.416 | 12% |
| DMR-029 | *A. humilis* | Director's Bay | 12.7 | 0.431 | 12% |
| DMR-047 | *A. humilis* | Snake Bay | 9.6 | 0.436 | 9% |
| DMR-049 | *A. humilis* | Snake Bay | 6.6 | 0.454 | 9% |
| DMR-051 | *A. humilis* | Snake Bay | 5.9 | 0.458 | 8% |
| DMR-052 | *A. humilis* | Snake Bay | 4.7 | 0.498 | 8% |
| DMR-053 | *A. humilis* | Snake Bay | 3.5 | 0.359 | 14% |
| DMR-062 | *A. humilis* | Snake Bay | 5.3 | 0.408 | 10% |
| DMR-003 | *A. lamarcki* | Director's Bay | 45.3 | 0.405 | 7% |
| DMR-004 | *A. lamarcki* | Director's Bay | 46.8 | 0.427 | 7% |
| DMR-005* | *A. lamarcki* | Director's Bay | 46.9 | 0.466 | 9% |
| DMR-006 | *A. lamarcki* | Director's Bay | 6.9 | 0.460 | 18% |
| DMR-008 | *A. lamarcki* | Director's Bay | 39.2 | 0.508 | 18% |
| DMR-009 | *A. lamarcki* | Director's Bay | 40.9 | 0.496 | 15% |
| DMR-010* | *A. lamarcki* | Director's Bay | 41.7 | 0.593 | 9% |
| DMR-011 | *A. lamarcki* | Director's Bay | 8.3 | 0.449 | 11% |
| DMR-014 | *A. lamarcki* | Director's Bay | 11.5 | 0.523 | 13% |
| DMR-016 | *A. lamarcki* | Director's Bay | 13.4 | 0.405 | 8% |
| DMR-017 | *A. lamarcki* | Director's Bay | 14.5 | 0.396 | 10% |
| DMR-018 | *A. lamarcki* | Director's Bay | 17.8 | 0.423 | 6% |
| DMR-019 | *A. lamarcki* | Director's Bay | 19.8 | 0.496 | 12% |
| DMR-020 | *A. lamarcki* | Director's Bay | 21.5 | 0.435 | 8% |
| DMR-021 | *A. lamarcki* | Director's Bay | 25.2 | 0.452 | 6% |
| DMR-023 | *A. lamarcki* | Director's Bay | 35.9 | 0.480 | 7% |
| DMR-024 | *A. lamarcki* | Director's Bay | 37.5 | 0.497 | 5% |
| DMR-025 | *A. lamarcki* | Director's Bay | 29.4 | 0.422 | 6% |
| DMR-026 | *A. lamarcki* | Director's Bay | 9.7 | 0.416 | 8% |

**Table 3.** Genome size estimates of *Agaricia* samples (continuation of previous page)

| Lab ID | Morphospecies | Collection site | Depth (m) | Genome size (pg) | CV |
| --- | --- | --- | --- | --- | --- |
| DMR-028 | *A. lamarcki* | Director's Bay | 10.3 | 0.438 | 8% |
| DMR-30 | *A. lamarcki* | Director's Bay | 15.8 | 0.450 | 11% |
| DMR-31 | *A. lamarcki* | Director's Bay | 18.5 | 0.370 | 10% |
| DMR-032 | *A. lamarcki* | Director's Bay | 19.9 | 0.419 | 16% |
| DMR-034 | *A. lamarcki* | Director's Bay | 35.3 | 0.407 | 11% |
| DMR-035 | *A. lamarcki* | Director's Bay | 39.1 | 0.546 | 9% |
| DMR-038 | *A. lamarcki* | Snake Bay | 31.8 | 0.453 | 8% |
| DMR-041 | *A. lamarcki* | Snake Bay | 29.1 | 0.440 | 13% |
| DMR-042 | *A. lamarcki* | Snake Bay | 26.3 | 0.423 | 8% |
| DMR-043 | *A. lamarcki* | Snake Bay | 22 | 0.394 | 10% |
| DMR-044 | *A. lamarcki* | Snake Bay | 16.7 | 0.407 | 9% |
| DMR-045 | *A. lamarcki* | Snake Bay | 15.6 | 0.442 | 7% |
| DMR-056 | *A. lamarcki* | Snake Bay | 32.3 | 0.411 | 10% |
| DMR-057 | *A. lamarcki* | Snake Bay | 26.7 | 0.439 | 8% |
| DMR-068 | *A. lamarcki* | Snake Bay | 33.5 | 0.439 | 12% |

## Notes

### Competing Interest Statement

The authors have declared no competing interest.

